# Is mild ADHD beneficial? Brain criticality peaks at intermediate ADHD symptom scores across the population

**DOI:** 10.1101/2022.12.14.519751

**Authors:** Hamed Haque, Jonni Hirvonen, Sheng H. Wang, Jaana Simola, Isabel Morales-Muñoz, Benjamin U. Cowley, J. Matias Palva, Satu Palva

## Abstract

Attention-deficit/hyperactivity disorder (ADHD) is characterized by a continuum of symptoms including inattentiveness, hyperfocus, and hyperactivity, that are manifested, *e.g.*, in increased reaction-time variability in continuous performance tasks (CPTs). Framework of brain criticality posits that brains operate in an extended regime of critical-like dynamics where neuronal and behavioral processes exhibit scale-free long-range temporal correlations (LRTCs) across hundreds of seconds. Whether shifts across critical-like brain states and parallel changes in LRTCs underlie ADHD symptoms has remained unexplored. We investigated whether brain-state shifts towards excitation-dominated dynamics and associated changes in LRTCs could explain ADHD symptoms and their continuum across individuals. We measured brain activity with magnetoencephalography (MEG) during resting and two CPTs from adult participants diagnosed with ADHD (*N* = 34) and neurotypical controls (NC) (*N* = 36) and characterized criticality with LRTCs of neuronal oscillations. ADHD patients exhibited stronger LRTCs than NC in low (5-20 Hz) and high (30-100 Hz) frequencies and dichotomous task effects. High-frequency LRTCs were positively correlated with ADHD symptoms, while beta-band (20-30 Hz) activity exhibited a quadratic correlation, peaking at moderate scores across the ADHD and NC cohorts being smaller for low and high scores. We demonstrate that ADHD is associated with shifts in brain criticality in resting-and task-state brain dynamics. Individuals with intermediate symptom scores across the NC and ADHD cohorts operated at peak criticality that is associated with a several functional benefits The progressive shift towards the supercritical side of the critical regime becomes detrimental only at moderate and severe symptoms.

## Introduction

Approaches from complex-systems and statistical-physics research have opened new avenues for understanding emergent brain dynamics and how they may contribute to the functional deficits in brain disorders. Attention-deficit/hyperactivity disorder (ADHD) is a heritable neurodevelopmental condition characterized by detrimental levels of impulsive, inattentive and/or hyperactive traits (1), as well as hyper-focus (2) and novelty seeking (3). Cognitive dysfunction in individuals with ADHD is reflected in increased reaction time variability (RTV) (4,5), which is thought to reflect intermittent drifting of attention and failures in executive functions (6). These excessive functional fluctuations are especially observable in continuous performance tasks (CPTs) that require sustained attention over long periods of time in a monotonous or repetitive environment. However, the underlying neuronal dynamics have remained poorly understood. Previous studies have established that alpha (α, 8−13 Hz) and beta (β, 20−30 Hz) band oscillations, which are fundamental for attention and top-down control (7–10), show reduced task-dependent suppression and lateralization of local oscillation amplitudes in adults (11–14) and children (15–17) with ADHD. However, ADHD is associated with the inability to sustain attention in timescales from seconds to minutes, which cannot be explained by sub-second timescale oscillation anomalies.

The brain criticality framework offers a statistical-physics mechanism for explaining variability, dynamics, and emergence in neuronal population activity (18,19). This framework thus links brain states and dynamic “operating points” of neuronal systems with brain-activity observables such as fluctuations and long-range correlations. The criticality framework posits that brains operate in an extended critical regime between subcritical and supercritical phases, *i.e.*, between disorder (asynchrony) and order (hypersynchrony) (20), respectively. In this regime, power-law spatio-temporal correlations emerge in brain dynamics and yield functional benefits, such as maximal dynamic range, information transmission, and representational capacity, which support optimal brain function (21,22). However, even healthy individuals express great variability in the individual operating point within the critical regime, which explains both inter-individual and-areal variability in mean levels of inter-areal synchronization (20). The primary control parameter responsible for tuning the operating-point is thought to be the net functional excitation/inhibition (E/I) ratio that is actively modulated by the ascending arousal system (AAD). Healthy brains appear to exhibit a slightly inhibition-dominated net E/I ratio, which poises them to operate on the subcritical side of the extended critical regime. Falling out of this regime through excessive inhibition or excitation leads to operation in the sub-or supercritical phase, which express either inadequate neuronal communication or epileptiform hyper-synchronized activity, respectively (20,23–28).

ADHD is associated with a shift in E/I towards greater excitability through both elevated glutamatergic excitation and GABAergic inhibitory deficits, especially in prefrontal cortex, basal ganglia and striatum (29,30). Importantly, several lines of evidence link this E/I shift also with AAS impairments and dysregulation of arousal-related neuromodulators, such as norepinephrine (NE) and dopamine, with ADHD and the executive function deficits therein (31). Conversely, the primary stimulant and non-stimulant medications for ADHD act on the NE and dopamine pathways of AAS to increase NE and dopamine availability in AAS projection targets (32–34). Excessive excitability in ADHD is further supported by shared risk factors (35) and notable co-morbidity between ADHD and epilepsy, where ADHD prevalence is greatly elevated in individuals with epilepsy and epilepsy prevalence and risk are elevated in individuals with ADHD (36).

We hypothesized that elevated cognitive variability in ADHD could be caused by these glutamatergic, GABAergic, and AAS impairments in E/I balance, such that they shift the operating point of the associated neuronal systems towards more excitation-dominated brain dynamics. In this model, this operating point shift translates the cellular-level and sub-second time-scale neurotransmission and modulation deficits into brain states that exhibit aberrant brain and behavioral dynamics in time scales from seconds to hundreds of seconds. As a hallmark observable of critical brain dynamics, neuronal oscillations exhibit long-range temporal correlations (LRTCs) in their amplitude envelopes lasting from seconds to minutes (19,37–39) and peaking at the idealized critical point (40–42). Inter-individual variability in neuronal LRTCs have been found to predict LRTCs in behavioral performance in CPT tasks, and, consequently, the lengths of behavioral streaks (37,38,43). LRTCs are the primary brain criticality construct that is directly associated with behavioral fluctuations. Previous studies show that LRTCs are attenuated in patients with epilepsy (44,45), Alzheimer’s disease (46), schizophrenia or schizoaffective disorders (47–49), and major depressive disorder (50,51), compared to neurotypical controls (NC) but elevated in patients with insomnia (52) and in the seizure onset areas in epilepsy patients (53). In patients with autism spectrum disorder, LRTCs have been shown to be both attenuated (54) and elevated (55,56). However, it has remained unclear whether ADHD symptoms are associated with altered LRTCs, despite that increased RTV, shifts in attentional states, and impulsivity in CPTs, indicate aberrant fluctuations in the 1-100 s time scales where there is most evidence for LRTCs in humans.

We thus tested here the hypothesis that ADHD symptoms could be explained in the brain criticality framework by a shift in individual operating points toward the super-critical side of the extended critical regime (Fig. 1). To test this, we recorded MEG data during two different CPTs from adult ADHD and neurotypical control (NC) participants and estimated LRTCs from MEG oscillation amplitude fluctuations using detrended fluctuation analysis (DFA).

**Figure 1.**
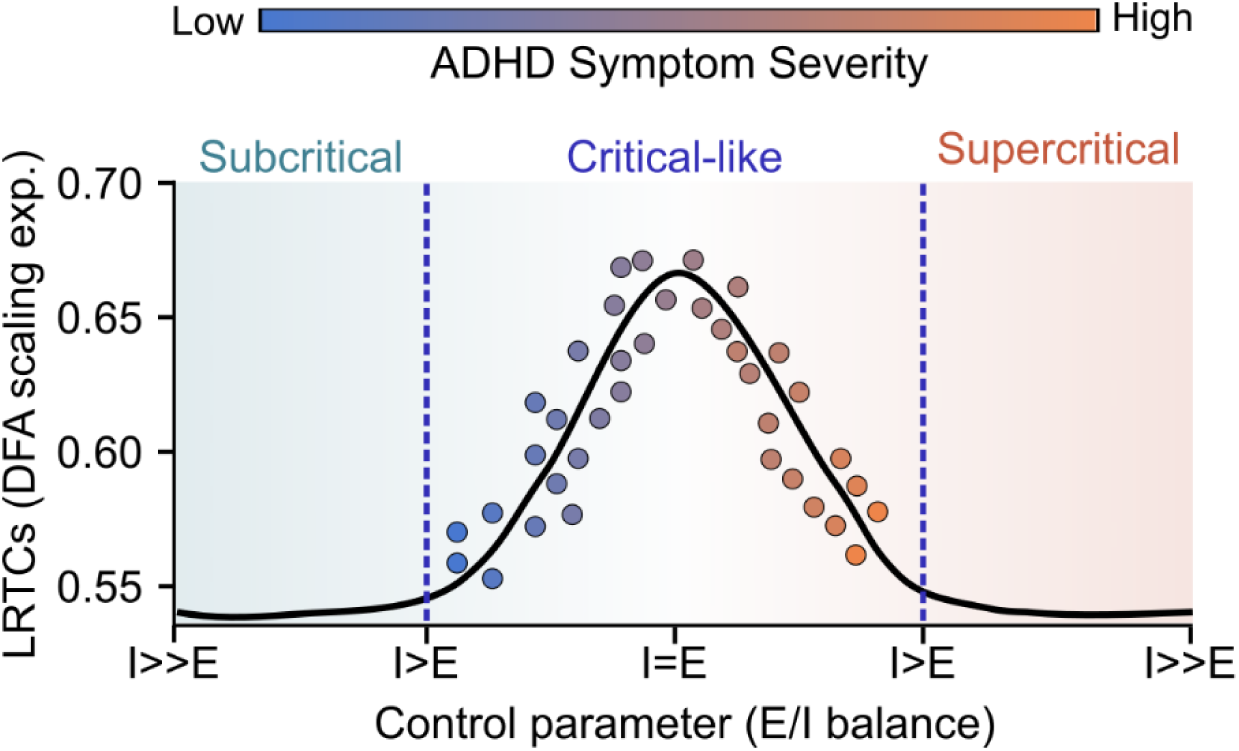
Schematic hypothesis of the role of brain criticality in ADHD brain mechanisms. We hypothesize that ADHD symptoms are predicted by an individual’s operating point in an extended critical regime. The individual’s operating point is determined by control parameters such as the excitation/inhibition (E/I) balance and AAS-based neuromodulation. While healthy human brains operate predominantly in the subcritical side of the critical regime, increase in ADHD symptom scores shifts the operating point closer towards the supercritical side of the critical regime.

## Methods and Materials

### Participants

The sample included 36 NC participants (mean ±SD: 34±9 years old, 13 females, 1 left-handed), and 34 participants with a prior ADHD diagnosis (mean ±SD: 37±9 years old, 19 females, 7 left-handed) recruited from a previous study (57), the University of Helsinki student body, and the local community through social media advertisement. There were no significant differences in age (Mann-Whitney U-test, *p* = 0.084) or gender (Chi-square test, *p* = 0.097) between the two cohorts. However, there were significantly more left-handed participants in the ADHD cohort (Chi-square test, *p* = 0.019).

The inclusion criteria for all participants were: aged 18–60 years, had normal or corrected to normal vision, and were compatible with MEG and MRI. Inclusion criteria for the ADHD cohort required an existing ADHD diagnosis, made within the Finnish healthcare system and based on the International Classification of Diseases (ICD-10), as evaluated by medical professionals. Exclusion criteria included any comorbid psychiatric or neurological disease, substance abuse, or medications affecting the central nervous system. ADHD participants refrained from ADHD-specific medications 24 hours before the MEG recordings. This study was conducted in compliance with the Declaration of Helsinki and approved by the Research Ethics Board at the Helsinki University Central Hospital. All participants gave written informed consent prior to participation.

### Psychopathology

ADHD symptoms were assessed in all participants using the Barratt Impulsiveness Scale (BIS) (58) and Adult ADHD Self-Report Scale (ASRS) (59); see Table S1 and Table S2. We first validated that the participants in the ADHD cohort exhibited clinical symptoms as measured by the two scales and that participants in the NC cohort had scores consistent for neurotypical populations (see Table S1). For the ADHD cohort, the mean and standard deviation (SD) were 81.10 ± 12.46 for BIS and 17.41 ± 3.75 for ASRS (Fig. 2E). For the NC cohort, total scores were 60.13 ± 9.79 for BIS and 9.71 ± 3.49 for ASRS, both of which were significantly lower than in the ADHD cohort (ASRS, Mann-Whitney U-test, *p* = 5.2E-11; BIS, Mann-Whitney U-test, *p* = 7.6E-9, Fig. 2E). Although these symptom scores are not officially part of any diagnosis criterium for ADHD and have been used mostly for screening purposes or evaluating the response for treatments, a total BIS sum of 72 or above should classify an individual as highly impulsive (60). In the case of ASRS, the maximum total score is 30 but for screening purposes certain questions weigh more than others. Three participants from the NC cohort fulfilled the screening criteria for ADHD and were excluded from all subsequent analyses expect for that shown in Fig. 6. Overall, these data show that ADHD symptoms in the NC cohort were mostly subclinical.

**Figure 2.**
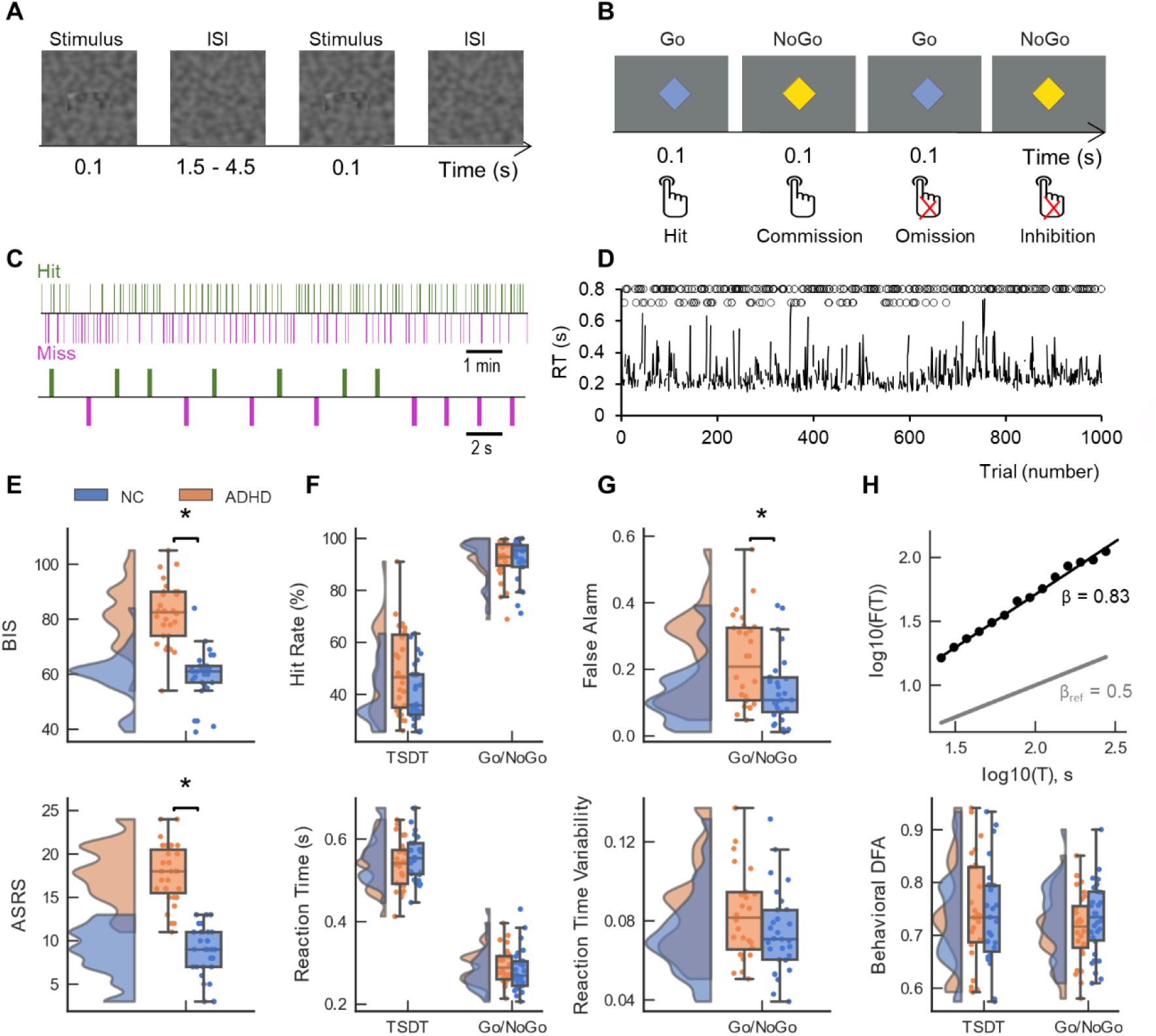
Task schematics and behavioral data. Schematics of (**A**) Threshold of Stimulus Detection Task (TSDT) and (**B**) Go/NoGo task experiments. (**C**) A short snapshot period of Hit-Miss time-series of a representative participant in TSDT. (**D**) Reaction Time (RT) series of a representative participant for each trial in Go/NoGo. Circles in the lower line above the time-series represent commission errors and circles in the upper line are trials with successful inhibition. (**E**) Distribution of BIS and ASRS scores for the ADHD (orange) and NC (blue) cohort. The box extends from the lower quartile (Q1, 25^th^ percentile) to the upper quartile (Q3, 75^th^ percentile). The line at the center of the boxplot indicates the median and the whispers represent the range of the data. The black bar above the boxplots with asterisk denotes significant differences between the ADHD and NC cohorts (Mann-Whitney U-test, *p* < 0.05). (**F**) Distribution of Hit rate (HR), and RT distributions separately for TSDT and Go/NoGo tasks. (**G**) Same as F but for FA rate and RTV (**H**) Top: the fitting for the behavioral DFA exponents in a representative subject from the ADHD cohort for the TSDT task. Black line indicates the fitting and the gray line the fitting for the null hypothesis. Bottom: behavioral DFA of the NC and ADHD cohorts (bottom).

### Tasks and Experiments

MEG data were collected during resting state (RS) and two visual CPTs: a Visual Threshold of Stimulus Detection Task (TSDT) and a Go/NoGo task. As reliable estimation of DFA exponents within an individual requires at least 10 minutes of data (37), we collected data over 25 minute runs for TSDT and 17 minute runs for Go/NoGo. Tasks were conducted on either one or two days, with the task order counterbalanced across participants. Four participants only performed the TSDT and four only performed the Go/NoGo task, leaving *N* = 62 total participants who performed both tasks. Both tasks were presented onto a projector screen inside a magnetically shielded room during MEG recordings. Participants provided responses via a button press (dual handheld RESPONSEPixx/MRI).

### Resting state

Before the task data collection, participants underwent 10 minutes of eyes-open resting state (RS) MEG recording, during which they fixated on a central cross. Data for RS were collected from *N* = 36 NC participants and *N* = 34 ADHD participants.

### Visual Threshold of Stimulus Detection Task (TSDT)

In the TSDT, participants detected visual stimuli presented at the perceptual threshold as described in J. M. Palva et al. (2013) (Fig. 2A). The visibility was individually pre-calibrated to achieve a 50% detection rate using the adaptive QUEST algorithm (Table S1). Here, prolonged periods or ‘streaks’ of consecutive misses reflect lapses in sustained attention. The task was programmed and executed using MATLAB (MathWorks, Natick, MA, USA). Visual stimuli of two shapes (50% probability) were presented in any orientation with duration of 0.1 seconds in slowly-varying gradient noise with a diameter of 10° (61) and with a random inter-stimulus interval (ISI) of 1.5–4.5 s. Each participant completed 500 stimulus trials over ∼25 minutes.

### Go/NoGo task

The Go/NoGo task was performed as in Simola et al. (2017), where participants responded as quickly as possible to Go stimuli (blue diamond) and refrained from responding to NoGo stimuli (yellow diamond) (Fig. 2B). Commission errors (false alarms) were erroneous responses to NoGo stimuli and constituted a measure of insufficient cognitive control. The task was coded using Presentation™ software (Neurobehavioral Systems, Inc., Albany, CA, USA). The stimulus was presented with a size of 1° for 0.1 s on a grey background (Fig. 2B). The Go (75% of trials) and NoGo (25% of trials) stimuli were distributed randomly in the stimulus stream with SOA of 1 s. Each participant completed 1000 trials over ∼17 minutes.

### Behavioral Data analysis

Hit Rates (HRs) for TDST were the proportion of correct detection for responses at latencies 200–1500 ms. Seven participants from the NC cohort and five participants from the ADHD cohort with HR < 25% were excluded from the analysis leaving *N* = 27 ADHD and *N* = 27 NC participants. In the Go/NoGo task, HRs were computed for the correctly responded Go-trials (‘Hits’) and incorrectly responded NoGo-trials (‘Commission errors’) for responses within 150–800 ms from stimulus onset. The false alarm rate was computed as the number of incorrectly responded NoGo-trials divided by the total number of NoGo trials presented. Five participants from both the NC and ADHD cohorts with HR < 65% for the Go stimuli were excluded leaving *N* = 28 ADHD and *N* = 28 NC participants. Reaction time variability (RTV) was derived by computing standard deviations over the individual RTs in Go-trials. Behavioral LRTCs were computed over Hit and Miss time-series in TSDT as in J. M. Palva et al. (2013) and in the Go/NoGo task for the RTs time-series as in Simola et al. (2017), using Detrended Fluctuation Analysis (DFA) as in (63).

### MEG and MRI Recordings

MEG data (Elekta Neuromag TRIUX) were recorded at the BioMag laboratory in the HUS Medical Imaging Centre with a 1000 Hz sampling rate and 0.03–330 Hz online passband. Electrooculography data were recorded for ocular artifact removal. T1-weighted anatomical MRI scans for cortical surface reconstruction were obtained from every participant at a resolution of 1 x 1 x 1 mm (MP-RAGE) with a 1.5 T MRI scanner (Siemens, Germany).

### MEG preprocessing, source analysis, and surface parcellations

Source reconstruction and data analysis followed previously published procedures (64–67). Maxfilter software (Elekta Neuromag) was used to suppress extra-cranial noise and interpolate bad channels. Independent component analysis (ICA) (MATLAB) was used for removing ocular and cardiac artefacts. MEG data were filtered using Morlet wavelets between 3–150 Hz with the time-frequency compromise parameter *m* = 5. Freesurfer (68) was used for volumetric segmentation of the MRI data, surface reconstruction, flattening, and creating the cortical parcellation using the Destrieux atlas. We used a 400-parcel parcellation, obtained by iteratively splitting the largest parcels of the Destrieux atlas (69) along their most elongated axis. This process applied the same parcel-wise splits for all subjects, resulting in roughly equal-sized parcels (70,71). Source modelling was performed with MNE using noise-covariance matrices with a regularization constant of 0.05 and computed from filtered (200–250 Hz) baseline data (-0.75–0.25 s). Source-space vertices were collapsed into the 400 parcels of the Destriux atlas using a fidelity-weighted collapse operator (72).

### Analysis of LRTCs from Neuronal MEG Data

LRTCs of oscillation amplitude fluctuations were estimated with DFA (19,37), which yields the scaling exponent *β.* This exponent is a measure of temporal clustering capable of predicting fluctuations in a one-dimensional time series over a long temporal range. *β* values between 0.5 and 1 indicate stronger temporal dependencies, which is an indication of criticality, while values closer to 0.5 are associated with uncorrelated noise. DFA exponents were computed per cortical parcel on amplitude envelopes of filtered parcel time-series, by segmenting data into time windows Δ*t* from 1–225 s (39,62,73,74). Each segment of integrated data was then locally fitted.

## Statistical analysis

Non-parametric Mann-Whitney U-test was used to test differences in behavioral performance between the NC and ADHD cohorts. The correlation between behavioral task performance with ASRS and BIS scores were assessed with Spearman’s ranked correlation test.

Before statistical testing, the 400 parcel MEG data were collapsed into 200 parcels to improve statistical stability by reducing the effects of inter-subject variability (66). Within-group differences between task-state and resting-state were obtained with Wilcoxon signed-ranked test separately for each frequency and parcel. Group differences were obtained using Welch’s *t-*test. Post-hoc analysis for the correlations between clinical scores and neuronal and behavioral scaling laws were estimated with Spearman rank correlation and quadratic correlations tests. Effect size estimates were obtained using Cohen’s *d* (75,76). Multiple comparisons were controlled with false discovery rate (FDR) correction by removing 5% of observed significant parcels (at α = 0.05). To remove remaining false positive observations, we estimated a threshold Q to define the proportion of significant observations that could thereafter arise by chance in any of the wavelet frequencies.

## Data/code availability statement

Raw electrophysiological data cannot be shared publicly due to regulations imposed by the Ethical Committees but can be shared for collaborative efforts upon request. All python code used in this work to produce results, run the statistical analyses, and create the final figures can be found at https://github.com/palvalab/ADHD.

## Results

We assessed behavioral measures using the TSDT (Fig. 2A) and Go/NoGo (Fig. 2B) tasks by analyzing the behavioral time-series data (Fig. 2C–D). As the TSDT task was pre-calibrated to a 50% stimulus-detection rate (Hit Rate, HR), as expected, the between-group differences in HRs were not significant for TSDT (Fig. 2F) (NC 40.26 ± 11.38% (mean ± SD), ADHD 48.8 ± 16.25%, Mann-Whitney U-test, *p* = 0.054) nor were there differences in reaction times (RTs) (NC 0.56 ± 0.058 s, ADHD 0.54 ± 0.059 s, Mann-Whitney U-test, *p* = 0.35). In the Go/NoGo task, neither HRs (NC 91.8 ± 8.17%, ADHD 91.9 ± 7.5%, Mann-Whitney U-test, *p* = 0.86), RTs (NC 0.28 ± 0.05 s, ADHD 0.29 ± 0.042 ms, Mann-Whitney U-test, *p* = 0.16), nor reaction time variability (RTV) (NC 0.074 ± 0.021 s, ADHD 0.084 ± 0.024 s, Mann-Whitney U-test, *p* = 0.098) were significantly different between the groups (Fig. 2F, G). However, the false alarm (FA) rate was larger for the ADHD cohort compared to the NC cohort (NC 0.14 ± 0.1, ADHD 0.22 ± 0.13, Mann-Whitney U-test, *p* = 0.018, Fig. 2G), as found previously (77,78). We then assessed LRTCs in the behavioral Hit Miss times-series (Fig. 2C) and RT time-series (Fig. 2D) data for the TSDT and Go/NoGo tasks, respectively, using DFA (Fig. 2H). DFA exponents did not differ between NC and ADHD cohorts for TSDT (0.74 ± 0.093 and 0.75 ± 0.1 respectively, Mann-Whitney U-test, *p* = 0.78, Fig. 2H), or the Go/NoGo task (0.73 ± 0.069 in the NC cohort and 0.71 ± 0.065 in the ADHD cohort, Mann-Whitney U-test, *p* = 0.34).

### Neuronal LRTCs are stronger in ADHD than in NC cohort

Oscillation amplitude envelopes were extracted from Morlet-wavelet filtered source-reconstructed MEG data (Fig. 3A) and DFA exponents were used to assess the LRTCs in the oscillation amplitudes (Fig. 3B). The mean DFA across all parcels peaked in the α-band (8−13 Hz) in both tasks and cohorts (Fig. 3C). Consistent with this, DFA exponents were smaller for NC than for the ADHD cohort also at the parcel-level in the α-band (8−13 Hz) in the TSDT task and in the gamma-band (γ, 35-65Hz) in the Go/NoGo task (Fig. 3D) (Welch’s *t*-test, *p* < 0.05, FDR corrected). To assess the robustness of these differences *post hoc*, we averaged the DFA exponents across the significant frequencies, which yielded significantly larger DFA exponents for ADHD than NC in both tasks (Fig. 3E) with moderate effect sizes of *d* = 0.54 for TSDT and *d* = 0.57 for Go/NoGo. Larger DFA exponents in the ADHD cohort were localized to the temporal and occipital cortices in TSDT, as well as in prefrontal regions and the precuneus (Fig. 3F). In Go/NoGo, differences in DFA exponents were observed in the occipito-temporal and visual cortices, and in the cingulate gyrus.

**Figure 3.** Differential modulations of neuronal DFA exponents in ADHD and NC cohorts at the whole-brain level. (**A**) Example of unfiltered neuronal time-series (top) and filtered oscillation amplitude time-series (bottom, gray trace) with the amplitude envelope (bottom, black trace) (**B**) DFA exponents extracted from oscillation amplitude envelope for a representative participant during TSDT (Threshold of Stimulus Detection Task) and Go/NoGo tasks. The scaling exponent β for 10 Hz and 30 Hz filtered data (black lines) and for uncorrelated noise (gray line). (**C**) Group-level mean DFA exponents averaged over all parcels as a function of frequency for ADHD (orange) and NC (blue) cohorts during both tasks. Shading shows 95% confidence intervals. (**D**) Group-level differences in the DFA exponents between NC and ADHD cohorts estimated separately for each cortical parcel (Welch *t*-test, *p* < 0.05, FDR corrected). The *y*-axis shows the fraction of cortical parcels (*P*) in which the difference was significant. DFA exponents were significantly greater in ADHD compared to NC in the α-band (8–13 Hz) for TSDT and in the γ-band (35–65 Hz) for Go/NoGo. (**E**) Distribution of individual DFA exponents averaged over significant frequencies and parcels from D for the α-band in TSDT and γ−band, for Go/NoGo task. Boxplots were drawn using the same convention as in Fig. 2. The bar shows statistical significance between NC and ADHD (two-tailed independent measures t-test, *p* < 0.05). (**F**) Cortical regions in which the differences in DFA exponents were observed in the α-band in TSDT and in the γ-band for Go/NoGo. The red color indicates the fraction of time-frequency elements (*P+*) that were significantly increased in ADHD compared to NC.

The α-band and γ-band effects also remained significant after controlling for age in the two cohorts using ANCOVA with age as a covariate (Fig. S1), indicating that the effects cannot be explained by age differences. In addition, we investigated whether differences in DFA exponents between the two cohorts may reflect differences in oscillation amplitudes between the two cohorts (Fig. S2). We found no significant differences in the mean parcel amplitudes between the NC and ADHD cohorts (Welch t-test, p < 0.05, FDR corrected), indicating that the DFA effects cannot be attributed to differences in signal-to-noise ratio.

### ADHD cohort shows different task-dependent modulation of LRTCs

Transitioning from a passive state (resting state) to an active task (TSDT or Go/NoGo) requires the engagement of attentional and executive networks. We thus investigated whether differences in DFA exponents between cohorts would emerge when comparing task-related activation compared to resting state (RS). In the NC cohort, α-band (8–13 Hz) DFA exponents were suppressed and γ-band (35−65 Hz) DFA exponents increased compared to the RS (Fig. 4A) (Wilcoxon signed rank test, *p* < 0.05, FDR corrected). However, in the ADHD cohort, there was only a slight suppression in the α-band DFA exponents in TSDT, and a robust increase in γ-band DFA exponents in Go/NoGo during task compared to RS (Fig. 4A).

**Figure 4.**
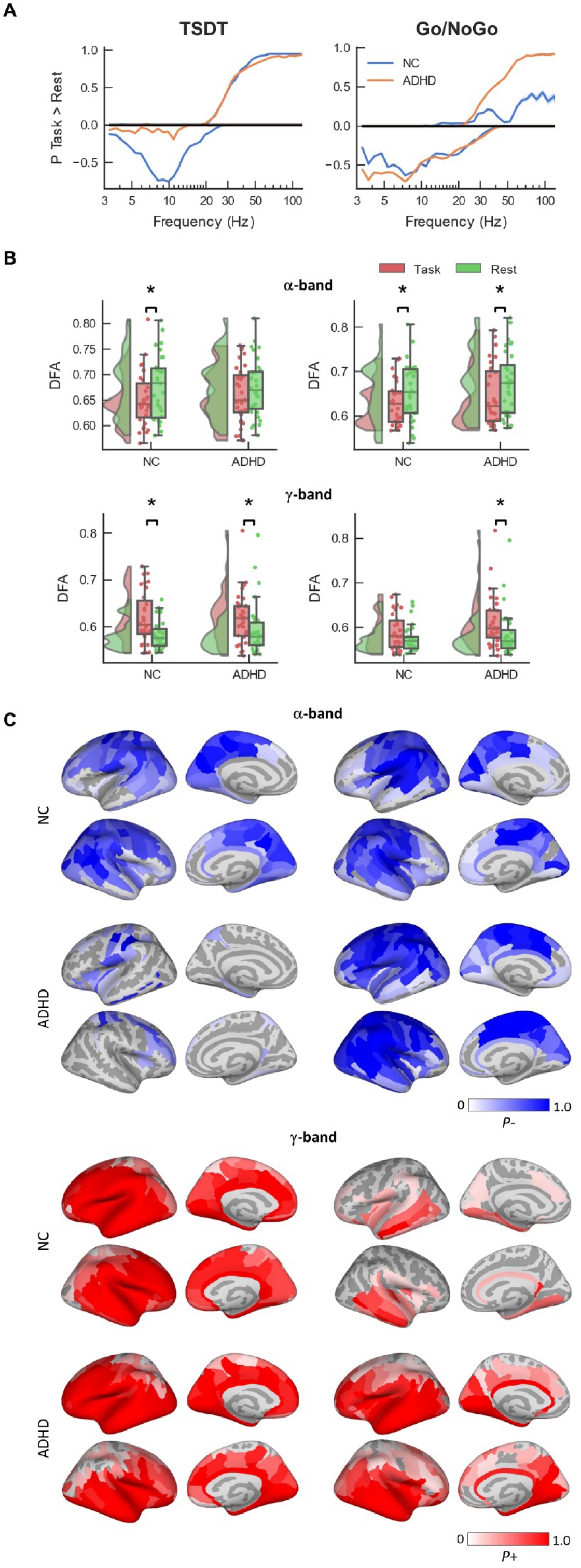
Task dependent modulations of neuronal DFA exponents. (**A**) Differences in DFA exponents between task and rest for the NC (blue) and ADHD (orange) cohorts (Wilcoxon signed rank test, *p* < 0.05, FDR corrected). Positive values indicate the proportion of cortical parcels where DFA exponents were larger during the Task compared to Rest (P+), while the negative values indicate the proportion of parcels where DFA exponents were larger during Rest than Task. Axis and shading as in Fig. 3D. (**B**) Distribution of individual DFA exponents for Task and Rest, separately for the NC and ADHD cohorts, in the α-band (8–13 Hz) (top) and γ-band (35–65 Hz) (bottom). Individual DFA exponents were computed from the average of all cortical parcels at the selected frequency range. The black bar with asterisk indicates significant differences in DFA exponents between Task and Rest (Wilcoxon signed-rank test, *p* < 0.01). (**C**) Cortical regions in which the DFA exponents were reduced in α-band by task performance (top) and increased in the γ-band by task performance (bottom). The red and blue colors indicate the fraction of time-frequency elements that were significantly increased (*P*+) or suppressed (*P*-), respectively.

To assess the reliability of the effects, we averaged individual DFA exponents in the α-band and γ-band for Task and Rest (Fig. 4B). The α-band DFA exponents were significantly higher during Rest than Task for the NC (moderate effect, *d* = 0.43) cohort in TSDT and both the NC (moderate effect, *d* = 0.42) and ADHD (moderate effect, *d* = 0.41) cohorts in Go/NoGo (Wilcoxon signed-rank test, *p* < 0.01). This difference was not significant for the ADHD cohort in the TSDT task (small effect, *d* = 0.23) For the γ-band, significant task modulation was observed for the ADHD cohort in TSDT (moderate effect, *d* =0.47) and Go/NoGo (moderate effect, *d* = 0.47), but for the NC cohort in only TSDT (large effect, *d* = 0.80). This difference was no significant for the NC cohort in Go/NoGo (small effect, *d* = 0.39).

The α-band suppression of DFA exponents was widespread in the NC cohort for both TSDT and Go/NoGo and was found predominantly in the visual and somatomotor systems (Fig. 4C). This pattern was also observed in the ADHD cohort for Go/NoGo but not for TSDT, where only a few frontal regions exhibited any significant task modulations. The increase in γ-band DFA exponents during Task was in nearly all brain regions for TSDT and in ventral visual cortex and the temporal regions for Go/NoGo.

### Neuronal LRTCs are correlated with ADHD symptoms

We then investigated whether these LRTC differences would explain ADHD symptoms, as quantified by ASRS and BIS symptom scales. We combined the data from both the NC and ADHD cohorts. DFA exponents in the high frequencies (30−100 Hz) were positively correlated with ASRS and BIS scores (Spearman’s rank correlation coefficient, *p* < 0.05, FDR corrected) (Fig. 5A), with greater number of significant parcels in Go/NoGo than TSDT. A post-hoc correlation test (Spearman’s rank correlation coefficient, *p* < 0.05) between the individual mean DFA exponents in the gamma frequency (47−52 Hz) and the ASRS and BIS scores revealed a significant correlation across individuals, this being robust for the Go/NoGo task (Fig. 5B).

**Figure 5.**
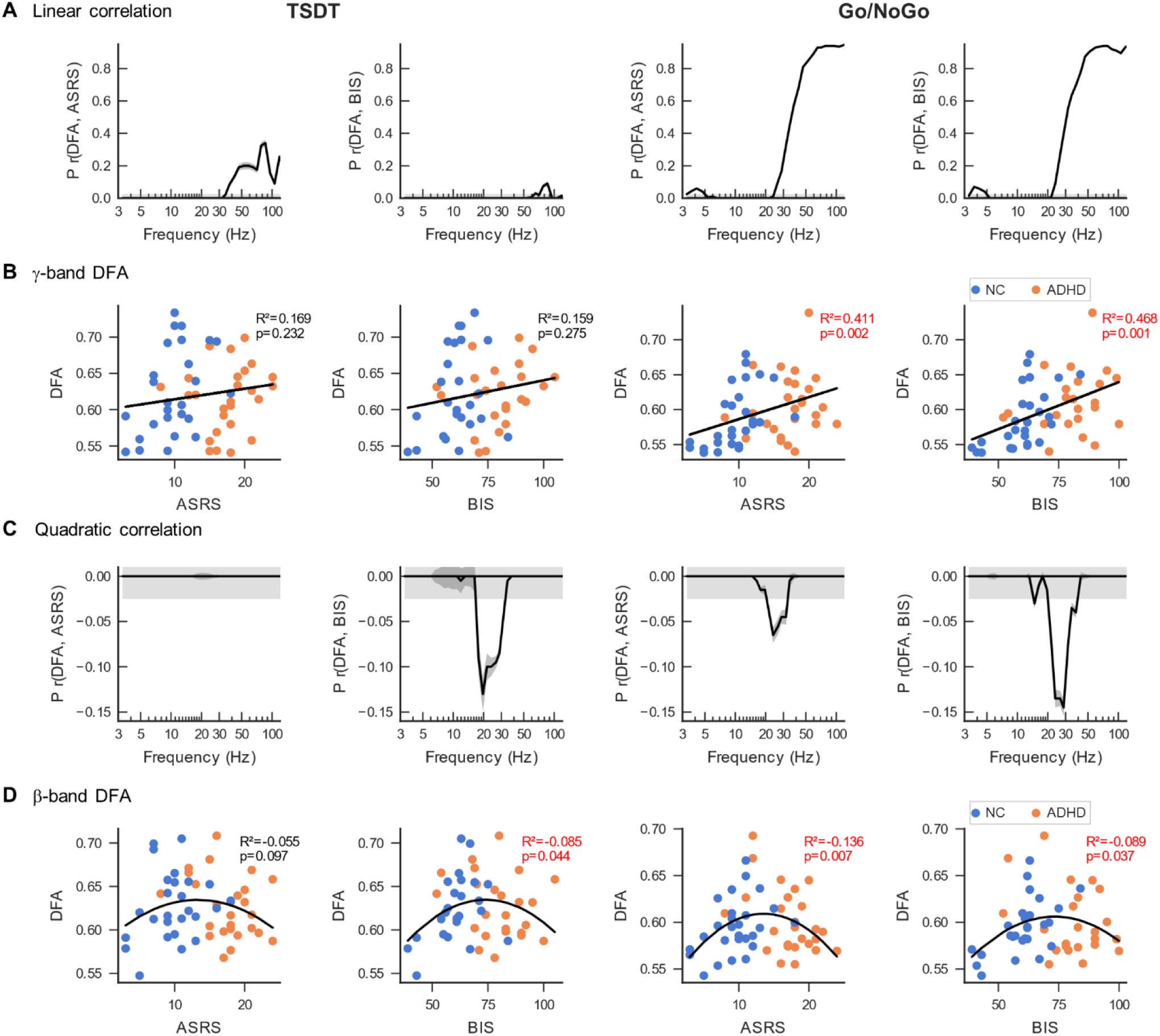
ADHD symptom scores predict neuronal LRTCs with an inverted U-shaped curve. (**A**) Linear correlation of neuronal DFA exponents obtained from task data with total ASRS and BIS scores (Spearman rank correlation test, *p* < 0.05, FDR corrected) for TSDT and Go/NoGo tasks. The *y*-axis shows the fraction of cortical parcels (*P*) where a significant correlation was observed. Participants from both the NC and ADHD cohorts were included. DFA exponents were significantly correlated with BIS and ASRS symptom scores in the γ-band. (**B**) Scatter plots for DFA exponents averaged over the γ-band (47−52 Hz) with BIS and ASRS scores. Data is plotted for the TSDT and the Go/NoGo tasks and include all participants from the ADHD (orange dots) and NC (blue dots) cohorts. (**C**) Same as in A but for the quadratic correlation between DFA exponents and ASRS or BIS scores. (**D**) Same as in B but for DFA exponents averaged over the β-band (22−24 Hz).

As LRTCs peak at the criticality, they could also have a quadratic relationship with symptoms if the symptoms are associated with operating point shifts across the peak of the critical regime. We estimated the quadratic DFA-symptom correlations for the combined NC-ADHD cohort with the linear component regressed out for all cortical parcels (Fig. 5C). In both the TSDT and Go/NoGo task, β-band DFA exponents were negatively quadratically correlated with ASRS and in the Go/NoGo also with BIS scores (Partial-quadratic correlation test, *p* < 0.05, FDR corrected). These quadratic correlations across subjects were confirmed with a post-hoc test (Partial-quadratic correlation test, *p* < 0.05) using individual mean DFA exponents averaged over all cortical parcels at a peak correlation frequency (22−24 Hz) (Fig. 5D).

## Discussion

LRTCs are a characteristic feature of brain activity (79), a hallmark of brain critical dynamics (42), and the primary brain-criticality correlate of behavioral fluctuations (37). Many brain disorders are associated with deviations from normative ranges of LRTCs (46–51,53–55), which are thought to reflect shifts in the operating points within an extended critical regime caused by shifts in E/I balance. Although ADHD is characterized by increased RTV on longer timescales in CPTs, deviations in LRTCs have not yet been established for ADHD. A single prior study has addressed whether ADHD was associated with altered LRTCs but found no evidence to this effect (80). Furthermore, current frameworks conceptualize mental disorders as a spectrum of symptoms rather than discrete diagnostic criteria, suggesting that deviations from normative dynamics should manifest as a continuum or as dimensions of dysfunction (81–83). Here, we advance the field in three fronts. First, we establish that ADHD-related deviations in LRTCs are due to differential rest-vs-task modulation. Second, we establish that these deviations arise in a continuum across symptom severity such that the strongest LRTCs were associated with moderate symptom scores, both in neurotypical individuals and those diagnosed with ADHD. Third, while several converging lines of evidence indicate a shift towards elevated excitability in ADHD, the mechanisms translating these factors into systems-level brain dynamics have remained unclear. We provide here evidence that this shift in E/I moves the operating point of brain dynamics in ADHD towards the supercritical side of the extended critical regime. Healthy brains operate in the subcritical side of the extended critical regime, while epileptogenic brain areas operate on its supercritical side (20). The present results provide a plausible explanation for the co-morbidity between ADHD and epilepsy through a shared shift of the operating point towards super-critical dynamics.

ADHD was associated with inadequate downregulation in the α-band during the TSDT task and an excessive upregulation in the γ-band during the Go/NoGo task. Previous studies have established sub-second time scales deviations in α-band oscillations in adults (11,12,14,84), adolescents (85–87), and children with ADHD (17,88,89) and in θ (θ, 4–8 Hz) (90,91) and γ-band (92,93) in adults. Here we show that ADHD is associated with deviant LRTCs in α-band and γ-band frequencies related to visual attention (7,10,94,95). Task differences may reflect the distinct cognitive demands whereby TSDT task requires sustained top-down visual attention, associated with tonic-like modulations of local (96,97) and inter-areal α-band oscillations (65,98), while the Go/NoGo task requires rapid and transient response inhibition and motor control linked to γ-band oscillations (99,100). This is supported by the localization of LRTCs to the prefrontal and occipital cortices associated with attention and executive functions (101,102) and in agreement with previous studies showing DAN (Dorsal Attention Network) hyperactivation and connectivity between DAN and VAN (Ventral Attention Network) in ADHD (103). Differences in LRTCs between ADHD and NC cohorts could also reflect prolonged brain maturation (104), which is a characteristic of ADHD (105,106).

Elevated LRTCs observed in ADHD are indicative of shifts in the brain’s operating point from the subcritical side towards more excitation-dominated super-critical states within the regime. As E/I balance and AAS modulation of E/I are key constructs for the biological control parameters determining this operating point, the shift observed in this study is congruent with several lines of evidence implicating excessive excitability and abnormal AAS regulation as the core biological basis of ADHD (29,32,33,107–109). Previous studies have considered the brain to operate within a single operating point in one individual brain state where brain disorders appear as a mean deviance from the optimal critical regime (22). We recently established that not only is there variability in individual operating points, but that there is also large variability across frequencies and brain anatomy whereby α and γ band oscillations operate closest to the critical point (20). Here we demonstrate the presence of frequency-specific deviances of individual operating points and their task-dependent changes in relation to ADHD symptoms and thus establish that brain disorders may arise via circuit-specific changes in brain critical states. Moreover, the opposing downregulation of α and upregulation of γ band oscillations’ “criticality” demonstrate the flexible dynamic control of individual operating points by behavioral demands. The present findings offer a plausible mechanistic explanation for how these sub-second synaptic-and micro-circuit-level characteristics of ADHD may be translated into the excessive multi-second and-minute cognitive and behavioral fluctuations that are pervasive in ADHD psychopathology.

Importantly, we found inverted U-shaped correlations between LRTCs and ADHD symptoms, with peak LRTCs at intermediate symptom scores, between neurotypical features and mild ADHD symptoms. This novel finding supports emerging frameworks which suggest that mental health symptoms, here associated with ADHD, arise from a continuum of brain dysfunction (81). As a unique perspective, these findings demonstrate that optimal brain function, as defined by brain criticality, is attained at moderate symptoms, while conversely, both low symptom scores in neurotypicals and high scores in ADHD cohort were associated with smaller LRTCs. Moreover, the results suggests that ADHD symptoms and their severity could be due to continuous increases of excitation leading to shifts in the brain operating point along the parameter space. Operation at the near critical regime is associated with beneficial neuronal dynamics, such as moderate levels of neuronal synchronization dynamics (20,42) and storage capacity (40), as well as beneficial behavioral performance including fluid intelligence (62,110). Therefore these results suggest that moderate ADHD symptoms scores, whether in clinical or control cohorts, are maximally associated with these functional benefits of brain critical dynamics. In contrast, both low symptom scores in the NC cohort and high symptoms scores in ADHD cohort were associated with smaller LRTCs.

In conclusion, these findings show that mild ADHD is be associated with brain dynamics that yield optimal functioning in exploratory behaviors (111,112), which is well aligned with behavioral ADHD-associated traits such as hyper-focus on highly motivating tasks (2,113), novelty-seeking (3) and sensation-seeking (114), which are evolutionary advantageous and facilitate adaptation to changing environments.

## Acknowledgements and Disclosures

This study was supported by funding from the Sigrid Juselius foundation to S.P. and J.M.P., Ella and Georg Ehrnrooth Foundation to I.M., and the Academy of Finland (SA 1294761) to J.S.

We thank Anna Lampinen for collecting the neuropsychological data.

H.H., J.H, and S.H.W. were responsible for formal analysis. H.H, J.H., J.S, and I.M. were responsible for investigation. H.H, J.H., and S.P. were responsible for writing the original draft of the manuscript. All authors were responsible for review and editing of the manuscript. H.H and J.H. were responsible for visualization. H.H. and S.H.W. were responsible for validation. H.H. was responsible for data curation. J.S., I.M., J.M.P., and S.P. were responsible for funding acquisition. J.M.P. and S.P. were responsible for conceptualization, supervision, and project administration.

A previous version of this article was published as a preprint on bioRxiv: https://doi.org/10.1101/2022.12.14.519751

The authors report no financial interests or potential conflicts of interest.

