## Supplementary Materials for "Is mild ADHD beneficial? Brain criticality peaks at intermediate ADHD symptom scores across the population"

| Subjects | Handedness | Age | Gender | Years of Education | ASRS Scores | BIS Scores | TSDT Threshold |
|---|---|---|---|---|---|---|---|
| N Total | N Left-handed | Mean±SD | N Females | Mean±SD | Mean±SD | Mean±SD | Mean±SD |
| **36 NC** | **1** | **33.81±9.27** | **13** | **18.40±3.84** | **9.71±3.49** | **60.13±9.79** | **0.025±0.013** |
| S0001 | right | 29 | Male | 20 | 8 | 60 | 0.0121 |
| S0002 | right | 29 | Male | 20 | 9 | 62 | 0.0104 |
| S0003 | right | 33 | Male | 22 | 10 | 62 | 0.0166 |
| S0004 | right | 41 | Female | 15 | 7 | 41 | N/A |
| S0005 | right | 33 | Female | 22 | 8 | 60 | 0.0259 |
| S0006 | right | 26 | Male | 19 | 11 | 62 | 0.0204 |
| S0007 | right | 26 | Male | 17 | 13 | 62 | 0.0133 |
| S0008 | right | 33 | Female | 23 | 9 | 61 | 0.0132 |
| S0009 | right | 26 | Male | 17.5 | 11 | 63 | 0.0175 |
| S0010 | right | 27 | Male | 9 | 13 | 84 | 0.0188 |
| S0011 | right | 33 | Male | 20 | 7 | 54 | 0.0204 |
| S0012 | right | 23 | Female | 17 | 5 | 43 | 0.0136 |
| S0013 | right | 26 | Female | 16 | 3 | 39 | 0.0229 |
| S0014 | right | 51 | Male | N/A | 7 | 67 | 0.0188 |
| S0015 | right | 18 | Male | 12 | 9 | 60 | 0.0182 |
| S0016 | right | 47 | Male | 30 | 3 | 43 | 0.0231 |
| S0017 | right | 23 | Male | 18 | 9 | 57 | 0.0192 |
| S0018 | left | 42 | Male | 12 | 12 | 72 | 0.0221 |
| S0019 | right | 34 | Male | N/A | N/A | N/A | 0.0278 |
| S0020 | right | 31 | Female | N/A | N/A | N/A | 0.0156 |
| S0021 | right | 27 | Female | N/A | N/A | N/A | 0.019 |
| S0022 | right | 57 | Female | 18 | 11 | 63 | 0.0294 |
| S0023 | right | 50 | Male | 14 | 5 | 56 | 0.0213 |
| S0024 | right | 34 | Male | 20 | 6 | N/A | N/A |
| S0025 | right | 39 | Male | 17 | N/A | N/A | 0.013304 |
| S0026 | right | 39 | Male | N/A | N/A | N/A | N/A |
| S0027 | right | 53 | Female | 17 | 13 | 58 | N/A |
| S0028 | right | 26 | Male | 18 | 16 | 57 | 0.040642 |
| S0029 | right | 35 | Male | 19 | 18 | 71 | 0.069019 |
| S0030 | right | 30 | Female | 17 | 15 | 75 | 0.039262 |
| S0031 | right | 25 | Male | 19 | 10 | 55 | 0.037929 |
| S0032 | right | 32 | Female | 23 | 12 | 57 | 0.010446 |
| S0033 | right | 30 | Male | 19 | 11 | 61 | 0.052356 |
| S0034 | right | 27 | Male | 20 | 11 | 67 | 0.04207 |
| S0035 | right | 41 | Female | 19 | 9 | 63 | 0.036222 |
| S0036 | right | 41 | Female | 21 | 10 | 69 | 0.029105 |

**Table S1: Demographic information for all participants in the neurotypical control (NC) cohort**. Subject identities were anonymized to maintain confidentiality. Ages and years of education were self-reported and taken with respect to the time of the study. ASRS and BIS scores were obtained via the Adult ADHD Self-Report Scale (ASRS) and the with Barratt Impulsiveness Scale (BIS), respectively. TSDT Threshold refers to the strength of the stimuli used during the individual calibration during the TSDT task, with higher numbers indicating more salient visual stimuli.

| Subjects | Handedness | Age | Gender | Years of Education | ASRS Scores | BIS Scores | TSDT Threshold |
|---|---|---|---|---|---|---|---|
| N Total | N Left-handed | Mean±SD | N Females | Mean±SD | Mean±SD | Mean±SD | Mean±SD |
| **34 ADHD** | **7** | **37.24±9.022** | **19** | **17.55±4.53** | **17.41±3.75** | **81.10±12.46** | **0.032±0.015** |
| P0001 | left | 48 | Male | N/A | 11 | N/A | N/A |
| P0002 | left | 36 | Male | N/A | N/A | N/A | N/A |
| P0003 | right | 49 | Female | N/A | 15 | N/A | 0.0195 |
| P0004 | right | 59 | Female | N/A | 16 | N/A | 0.0272 |
| P0005 | right | 30 | Male | 14 | 15 | 68 | 0.0208 |
| P0006 | left | 55 | Male | 17 | 12 | 54 | 0.0167 |
| P0007 | right | 41 | Female | 18 | 17 | 83 | 0.0226 |
| P0008 | right | 30 | Female | 17 | 21 | 69 | 0.0199 |
| P0009 | right | 29 | Female | 14 | 19 | 90 | 0.0229 |
| P0010 | right | 35 | Male | 12 | 17 | 90 | 0.0288 |
| P0011 | right | 43 | Male | 10 | 15 | 74 | 0.0204 |
| P0012 | right | 30 | Male | 14 | 8 | 52 | 0.0173 |
| P0013 | left | 30 | Female | 19 | 18 | 92 | 0.0201 |
| P0014 | right | 34 | Male | 17 | 19 | 81 | 0.0141 |
| P0015 | right | 28 | Female | 18.5 | 16 | 80 | 0.0269 |
| P0016 | right | 30 | Female | 19 | 17 | 78 | 0.0186 |
| P0017 | right | 28 | Male | 20 | 13 | 70 | 0.0323 |
| P0018 | right | 51 | Female | 19 | 22 | 90 | 0.0208 |
| P0019 | left | 29 | Female | 19 | 12 | 69 | 0.0411 |
| P0020 | right | 26 | Female | N/A | N/A | N/A | 0.0169 |
| P0021 | right | 38 | Female | 14 | 18 | 83 | 0.0242 |
| P0022 | right | 27 | Female | 28 | 16 | 85 | 0.0206 |
| P0023 | right | 29 | Male | 18 | 24 | 100 | 0.0162 |
| P0024 | left | 34 | Male | 20.5 | 21 | 89 | 0.045601 |
| P0025 | right | 51 | Male | 10 | 24 | 105 | 0.066676 |
| P0026 | right | 38 | Male | 17 | 20 | 78 | 0.071445 |
| P0027 | left | 47 | Female | 18 | 18 | 83 | 0.051165 |
| P0028 | right | 44 | Female | 17 | 21 | 74 | 0.044563 |
| P0029 | right | 32 | Female | 18 | 21 | 99 | 0.040176 |
| P0030 | right | 31 | Male | 18 | 21 | 82 | 0.05 |
| P0031 | right | 31 | Male | 15.5 | 18 | 95 | 0.044052 |
| P0032 | right | 45 | Female | 12 | 14 | 79 | 0.051757 |
| P0033 | right | 41 | Female | 27 | 20 | 89 | 0.046129 |
| P0034 | right | 37 | Female | 28.5 | 18 | 71 | 0.034592 |

**Table S2: Demographic information for all participants in the ADHD cohort. Details reported are same as in Table S1**.

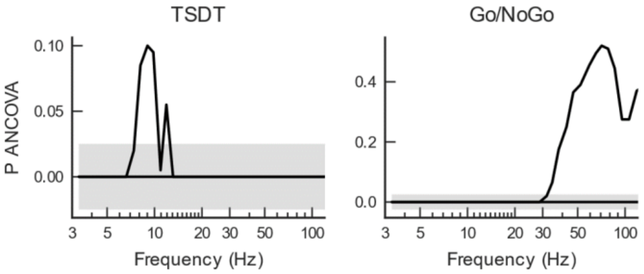


**Figure S1: The main effect of cohort (NC or ADHD) on DFA exponents, while controlling the effect of age**. An ANCOVA was conducted to control the effect of age in the differences of DFA exponents between the two cohorts. After controlling for age, there was a significant effect of group on DFA exponents in the α-band for TSDT and in the g-band for Go/NoGo. These effects of DFA exponents are in the same bands reported in Fig. 3D and therefore reveal that such cohort differences arise even when accounting for the age of the participants.

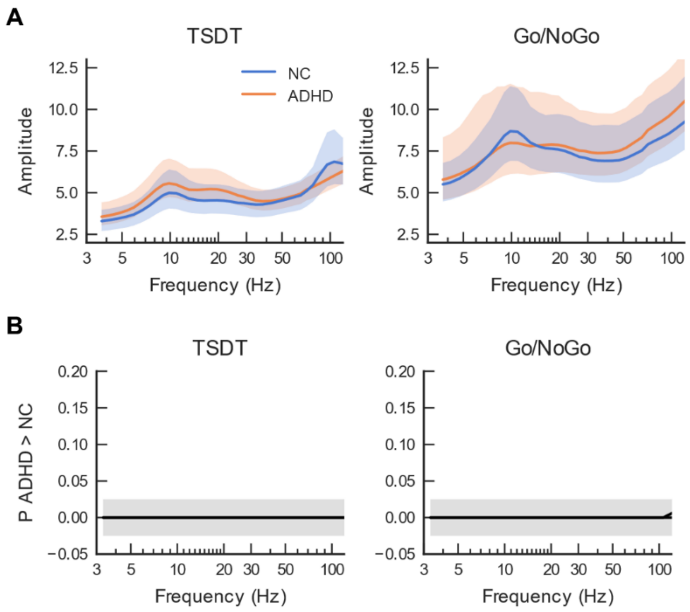


**Figure S2: Differences in DFA exponents cannot be explained by parcel amplitudes. (A)** Parcel amplitudes was computed for both Go/NoGo and TSDT for both the NC and ADHD cohorts. To compare the fluctuations of amplitude with neuronal LRTCs, parcel amplitudes were computed in continuous 4 s windows throughout the task and averaged. **(B)** The comparison between the parcel amplitudes in the two cohorts showed not significant differences (Welch t-test, $p < 0.05$, FDR corrected).
